## Supplementary Information for "The translation inhibitors kasugamycin, edeine and GE81112 target distinct steps during 30S initiation complex formation"

### Supplementary Figures

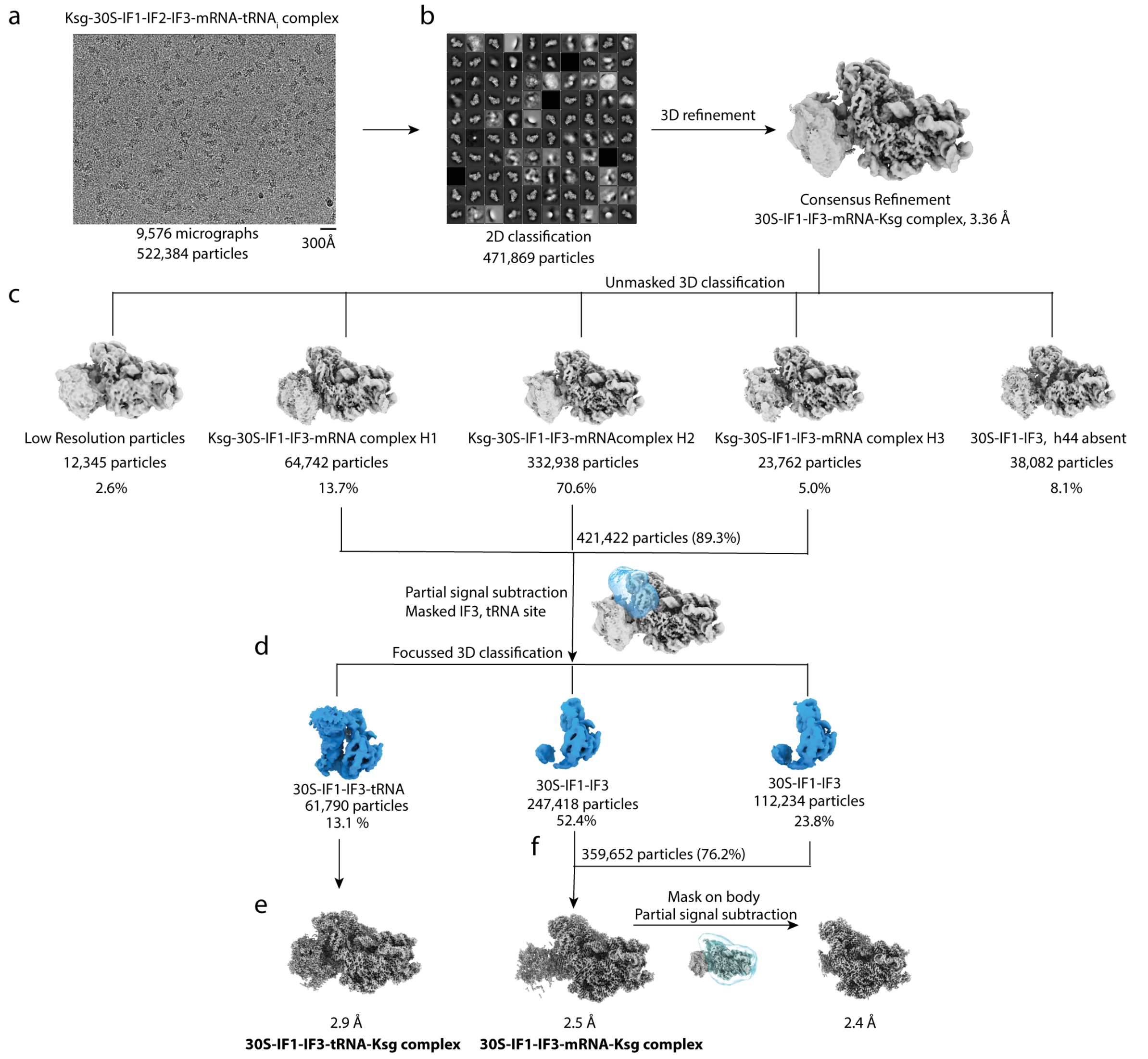


**Supplementary Fig. 1 | *In silico* sorting scheme of *E. coli* Ksg-30S complex. a,** From 9,576 micrographs, a total of 522,384 particles were picked by crYOLO using general model and subjected to 2D classification. **b,** After 2D classification, 471,869 particles were taken for initial consensus refinement. **c,** Unmasked 3D classification into 5 classes was performed. Three classes with Ksg density and different head movement (H1,H2,H3) were merged resulting in 421,422 particles (89.3%). **d,** Another round of focussed 3D classification was performed with a mask surrounding IF3 and tRNA site, yielding three classes, one with density for tRNA (13.1%, 61,790 particles) and other two with no density for tRNA present. **e,** The class with tRNA present (13,1%, 61,790 particles) was refined to high resolution to obtain 2.9 Å resolution. **f,** The latter two classed from (d) without density for tRNA was combined, resulting in 359,652 particles (76.2%). This was further refined to high resolution to reach 2.4 Å resolution. The 30S-head was removed by signal subtraction using a mask on the 30S-body.

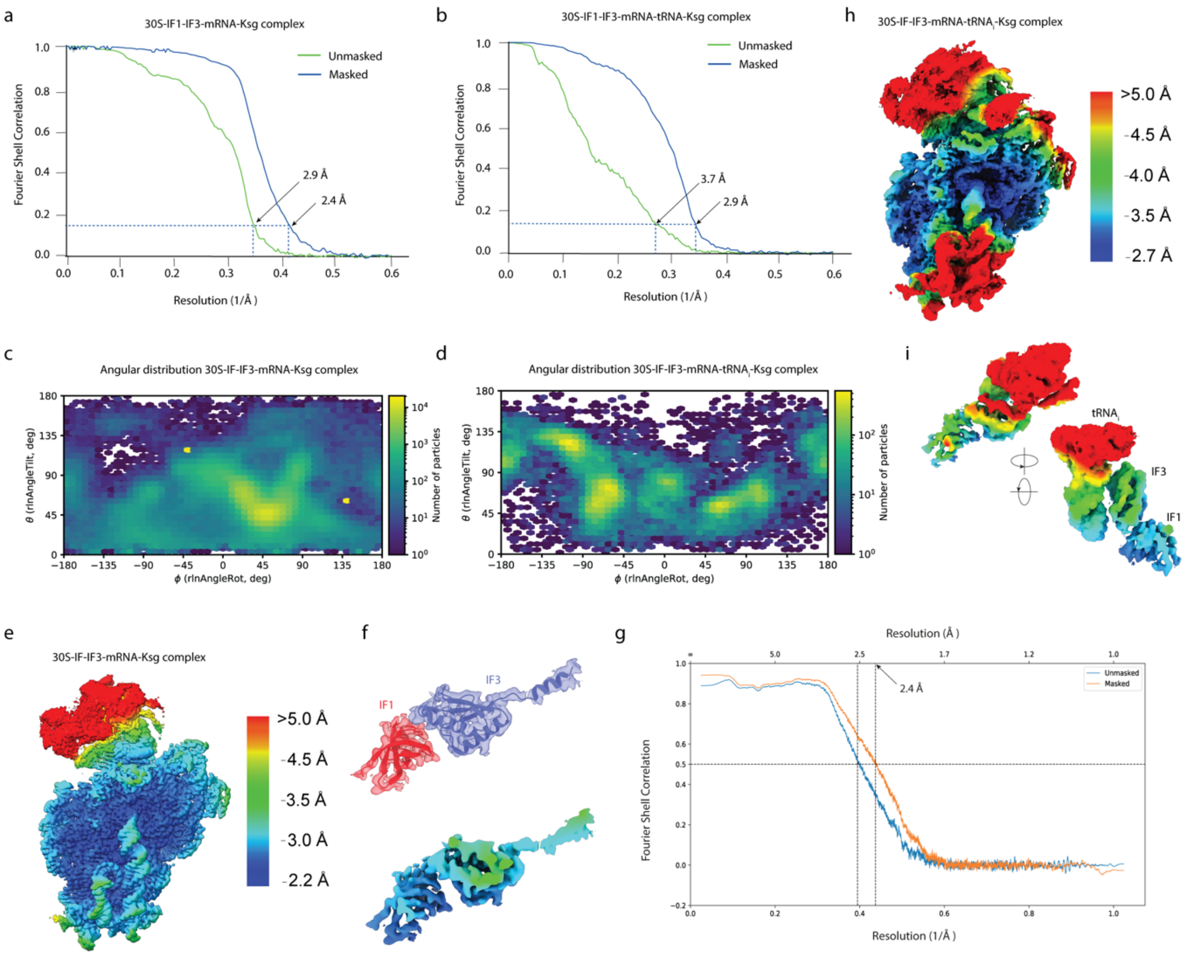


**Supplementary Fig. 2 | FSC and local resolution for *E. coli* Ksg-30S complex. a,** FSC curve for the Ksg-30S-IF1-IF3-mRNA complex map. **b,** FSC curve for the Ksg-30S-IF1-IF3-mRNA-tRNA complex map. **c**, Angular distribution plot for the Ksg-30S-IF1-IF3-mRNA complex map. **d**, Angular distribution plot for the Ksg-30S-IF1-IF3-mRNA-tRNA complex map. **e**, Overview of local resolution of Ksg-30S-IF1-IF3-mRNA complex map. **f**, Isolated densities with fitted models for IF1 and IF3 from map in (e), also colored according to local resolution. **g**, FSC map versus model for Ksg-30S-IF1-IF3-mRNA complex map. **h**, Overview of local resolution of Ksg-30S-IF1-IF3-mRNA-tRNA complex map. **i,** Isolated densities of IF1, IF3, tRNA from map in (e), colored according to local resolution.

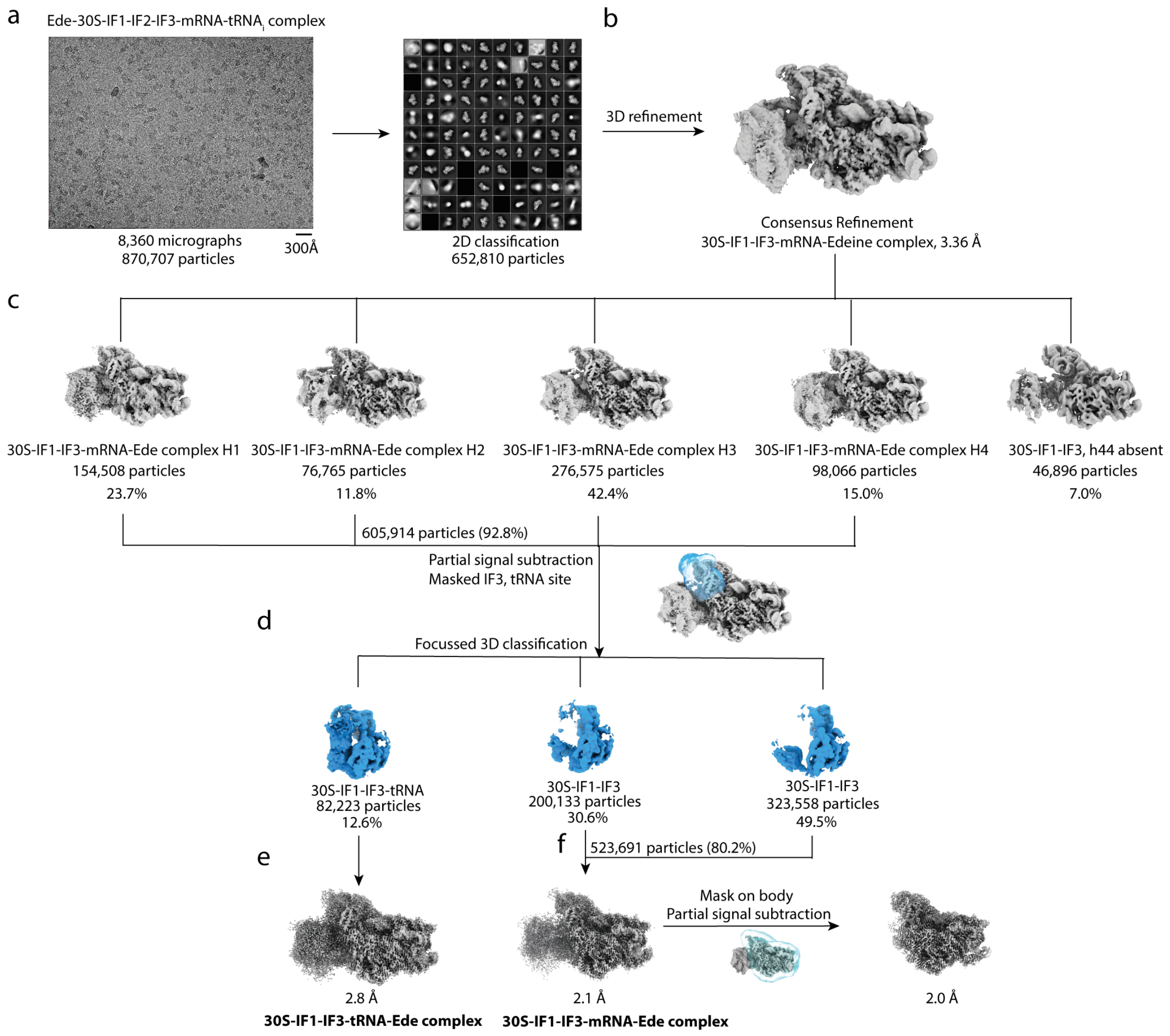


**Supplementary Fig. 3 | *In silico* sorting scheme of *E. coli* Ede-30S complex. a,** From 8,360 micrographs a total of 870,707 particles were picked by crYOLO using general model and subjected to 2D classification. **b,** After 2D classification, 652,810 particles were taken for initial consensus refinement. **c,** Unmasked 3D classification into 5 classes was performed. Four classes with Ede density and different head movement (H1,H2,H3, H4) were merged resulting in 605,914 particles (92.8%). **d,** Another round of focussed 3D classification after signal subtraction was performed with a mask surrounding IF3 and tRNA site, yielding three classes, one with density for tRNA (12.6 %, 82,223 particles) and other two with no density for tRNA present. **e**, The class with tRNA present (12.6%, 82,223 particles) was refined to high resolution to obtain 2.8 Å resolution. **f**, The latter two classed from d without density for tRNA was combined, resulting in 523,691 particles (80.2%). This was further refined to high resolution to reach 2.0 Å resolution. Head was removed by signal subtraction using a mask covering the 30S-body.

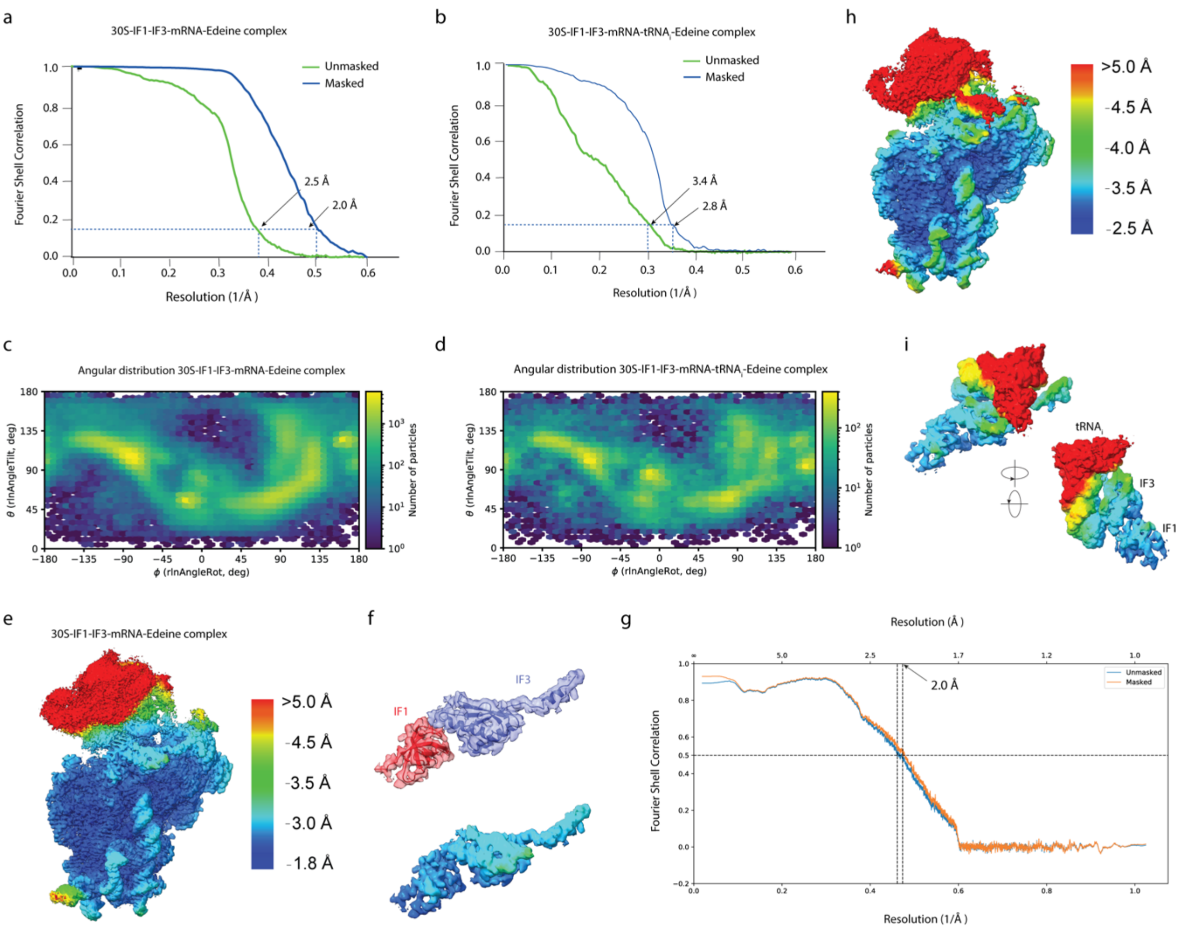


**Supplementary Fig. 4 | FSC and local resolution for *E. coli* Ede-30S complex. a,** FSC curve for the Ede-30S-IF1-IF3-mRNA complex map. **b,** FSC curve for the Ede-30S-IF1-IF3-mRNA-tRNA complex map. **c**, Angular distribution plot for the Ede-30S-IF1-IF3-mRNA complex map. **d**, Angular distribution plot for the Ede-30S-IF1-IF3-mRNA-tRNA complex map. **e**, Overview of local resolution of Ede-30S-IF1-IF3-mRNA complex map. **f**, Isolated densities with fitted models for IF1 and IF3 from map in e, also colored according to local resolution. **g**, FSC map versus model for Ede-30S-IF1-IF3-mRNA complex map. **h**, Overview of local resolution of Ede-30S-IF1-IF3-mRNA-tRNA complex map. **i,** Isolated densities of IF1, IF3, tRNA from map in (e), colored according to local resolution.

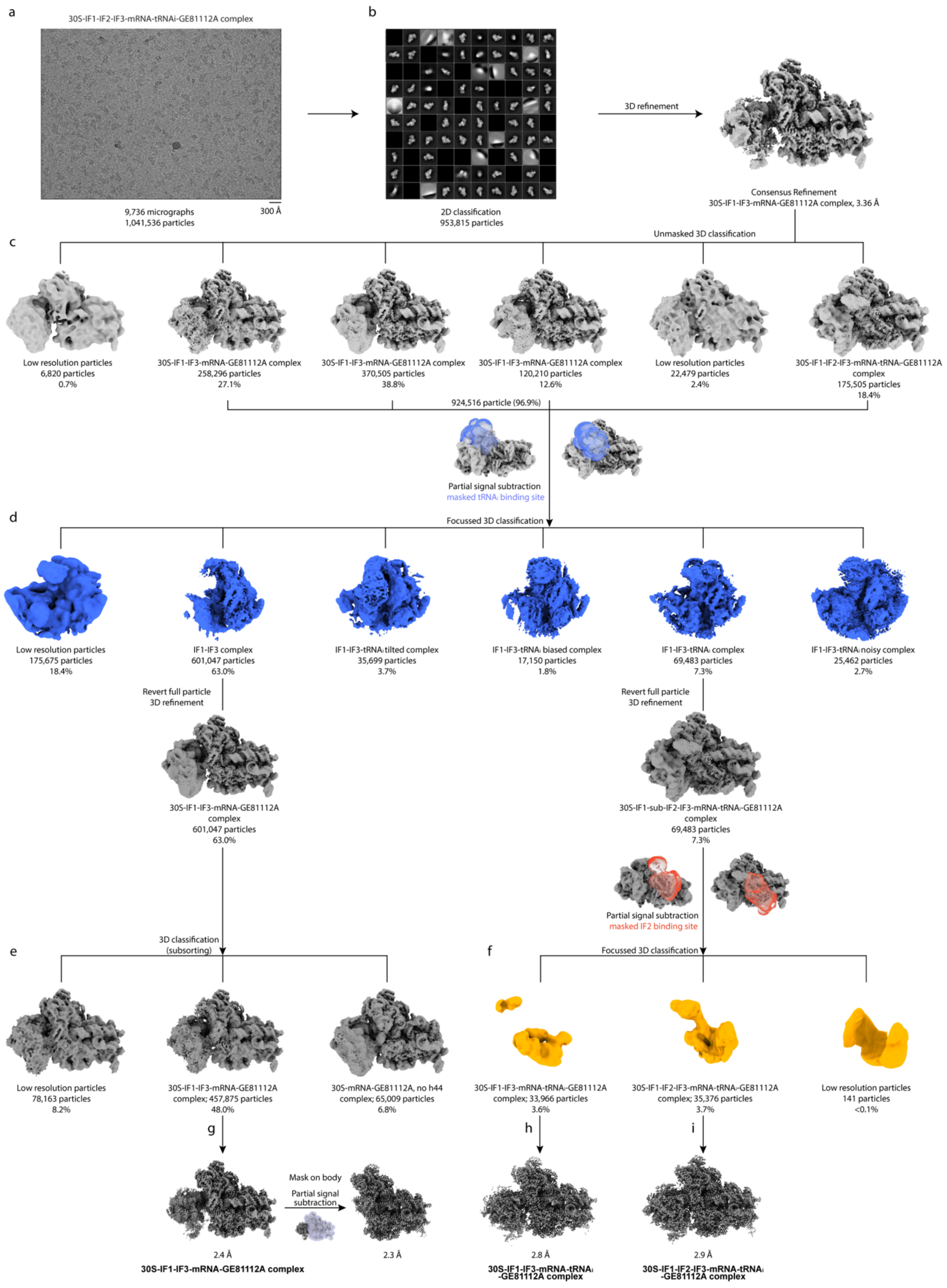


**Supplementary Fig. 5 | *In silico* sorting scheme of *E. coli* GE-30S complex. a,** From 9,736 micrographs, a total of 1,041,536 particles were picked by crYOLO using general model and subjected to 2D classification. **b,** After 2D classification, 953,815 particles were taken for initial consensus refinement. **c,** Unmasked 3D classification into 6 classes was performed. Four high resolution classes with Ede density were merged resulting in 924,516 particles (96.9%). **d,** Another round of focussed 3D classification after signal subtraction was performed with a mask surrounding tRNA site, yielding six classes, majorly one with density for tRNA (7.3 %, 69,843 particles) and another with no density for tRNA present (601,047 particles, 63%). **e**, The class without tRNA present was subsorted to remove junk particles **f**, Another round of focussed 3D classification was performed on tRNA classes after signal subtraction, with a mask surrounding IF2 site, yielding class with and without IF2. **g**, Class without tRNA was refined to 2.3 Å by masking the 30S body since the head was flexible. **h**, Class with tRNA but no IF2 and with IF2 were further refined to yielding 2.8 Å and 2.9 Å resolution, respectively.

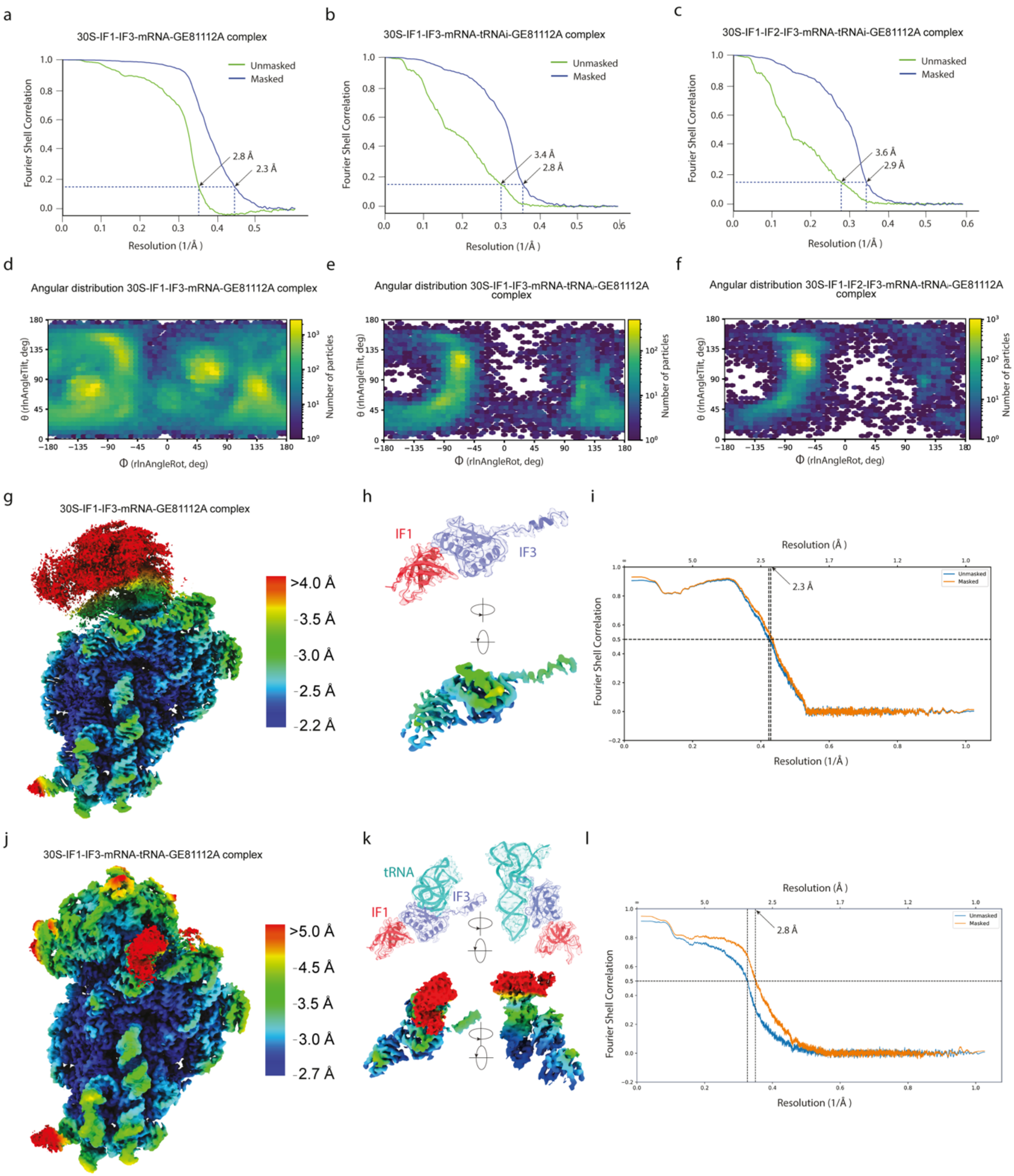


**Supplementary Fig. 6 | FSC and local resolution for *E. coli* GE-30S complex.** FSC curve for (**a**) GE-30S-IF1-IF3-mRNA complex map**,** (**b**) GE-30S-IF1-IF3-mRNA-tRNA complex map, (**c**) GE-30S-IF1-IF2-IF3-mRNA-tRNA complex map. Angular distribution of respective maps in (**d-f**). **g**, Overview of local resolution of GE-30S-IF1-IF3-mRNA complex map. **h**, Isolated densities with fitted models for IF1 and IF3 from map in (g), also colored according to local resolution. **i**, FSC map versus model for map in (g). **j**, Overview of local resolution of GE-30S-IF1-IF3-mRNA-tRNA complex map. **k,** Isolated densities with fitted models of IF1, IF3, tRNA from map in (j), colored according to local resolution. **l,** FSC map versus model for map in (j).

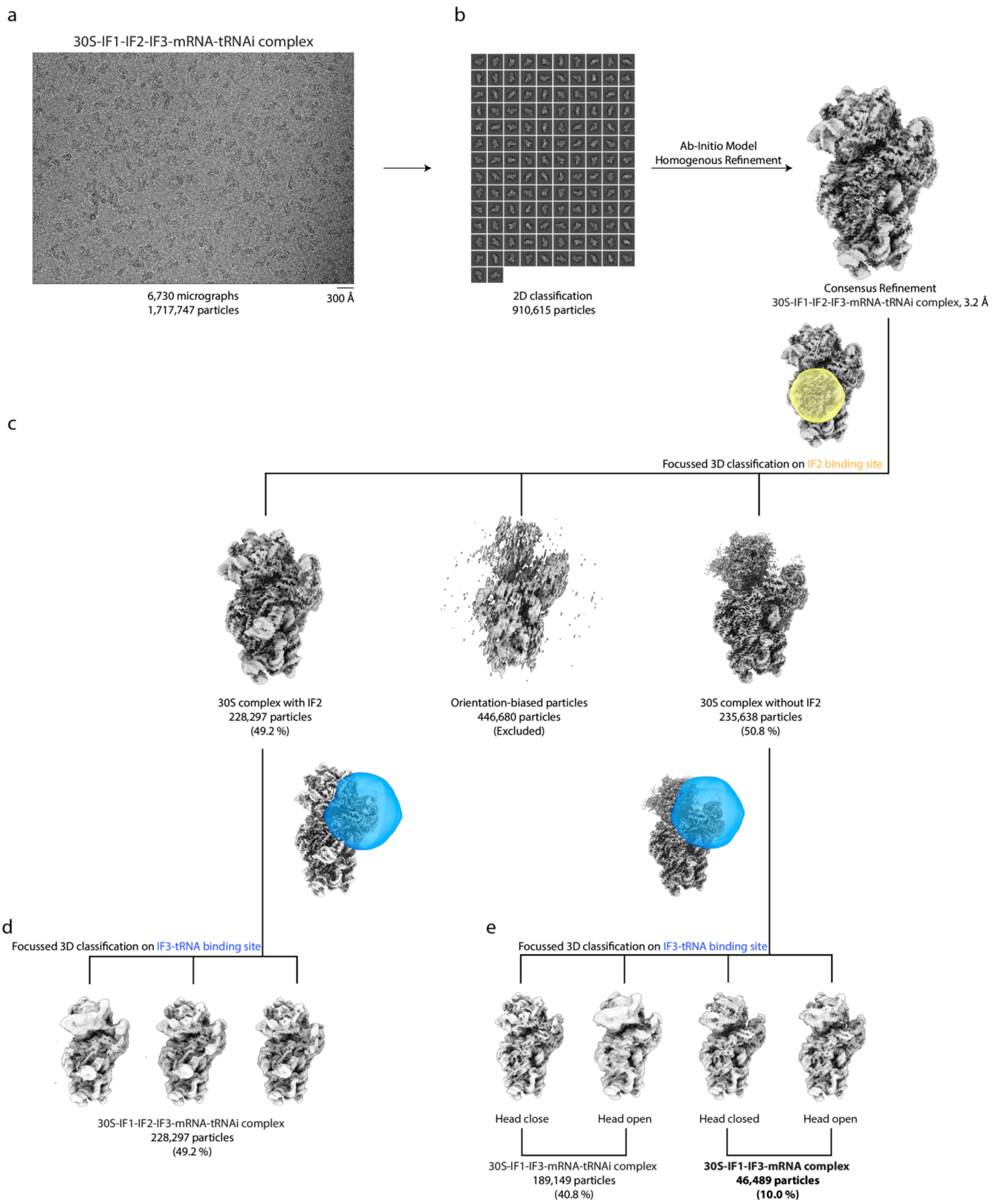


**Supplementary Fig. 7 | *In silico* sorting scheme of *E. coli* 30S initiation complexes in the absence of antibiotic. a,** From 6,730 micrographs, a total of 1,717,747 particles were picked by crYOLO using general model and subjected to 2D classification. **b,** After 2D classification, 910,615 particles were used for ab-initio model and homogenous refinement. **c,** Focussed 3D classification with mask around IF2 into 3 classes was performed. This resulted in class with IF2 (49.2%) and without IF2 (50.8%). Around 446,800 particles were discarded since they showed orientation bias. **d,** Another round of focussed 3D classification with mask around IF3 and tRNA binding site was performed on class with IF2 using a mask surrounding tRNA site, yielding three classes, with minor differences in IF3 conformation. **e**, Similarly, one round of focussed 3D classification with mask around IF3 and tRNA binding site was performed on class without IF2, yielding four classes. Two of them had tRNA density (189,149 particles, 40.8%) and two others were without tRNA (46,489 particles, 10%).
